# Activity of *Salmonella* SPI-1 inhibits the TLR4-dependent transcriptional but not translational response during macrophage infection

**DOI:** 10.1101/2024.01.08.574706

**Authors:** George Wood, Rebecca Johnson, Matt Brember, Filip Lastovka, Pani Tourlomousis, Clare Bryant, Betty Y-W Chung

## Abstract

Changes in gene expression during bacterial infection are the combined result of altered transcription and translation, with the latter comparatively understudied. Gram-negative bacteria rapidly trigger cytokine gene transcription in macrophages through the activation of pathogen associated molecular pattern receptors, for example detection of *Salmonella* lipopolysaccharide (LPS) from the bacterial cell envelope by Toll-like receptor 4 (TLR4). Here, through time-resolved parallel translatomic and transcriptomic profiling, we now show temporal TLR4-specific translational upregulation of cell signalling proteins in macrophages induced by *Salmonella*. While transcriptional upregulation of these genes is dampened through the activity of the *Salmonella* SPI-1 type three secretion system, a robust translational response remains. These data reveal an important host-pathogen translational regulatory network that modifies the innate immune response of macrophages to infection.

## Introduction

Rapid activation of the innate immune response is among the first lines of defence against invading pathogens and plays a crucial role in controlling and eliminating infection. Pathogen-associated molecular patterns (PAMPs) are detected by germ-line-encoded host receptors, triggering an innate immune response. Toll-like receptors (TLRs) detect a variety of PAMPs. This includes bacterial LPS, lipoproteins and flagellin, which are detected by TLR4, TLR2 (in combination with TLR1 or TLR6), and TLR5 respectively (Gay *et al*, 2014). Their activation in innate immune cells induces anti-microbial defences and triggers cytokine and chemokine production for recruitment and modulation of further immune cells to aid in fighting the infection (Satoh & Akira, 2016). As such, pathogen detection triggers dramatic alteration of host cell protein synthesis. The transcriptional response to bacterial pathogens is well studied, but our understanding of gene expression is incomplete without also examining translational regulation. While purified bacterial PAMPs have been shown to modulate translation of specific genes independently of transcription (Kitamura *et al*, 2008; Schott *et al*, 2014), there are few studies that dissects the TLR4-specific transcriptional response from the translational response to bacterial infection.

*Salmonella* are Gram-negative bacterial pathogens that can infect many different host cells and cause a range of diseases, from typhoid fever to gastroenteritis (Gal-Mor *et al*, 2014). On infection, *Salmonella* pivot to an intracellular lifecycle, supported by two type III secretion systems encoded on *Salmonella* pathogenicity islands (SPI) 1 and 2. SPI-1 is important for bacterial invasion of host epithelia, but macrophages can additionally take up bacteria by phagocytosis, whereas SPI-2 is expressed once the bacterium is intracellular to support *Salmonella* survival and replication (Wang *et al*, 2020). Both secretion systems also deliver effectors that modulate host cell immune signalling pathways (Galán, 2021; Sun *et al*, 2016; Jones *et al*, 2008; Pavlova *et al*, 2011; Jennings *et al*, 2017; Wang *et al*, 2020). TLR4 signalling is of key importance in *Salmonella* infection, where it plays roles in both infection control and bacterial virulence. Mice lacking TLR2 and TLR4 show increased mortality and bacterial burden in *Salmonella* infection (Weiss *et al*, 2004; Talbot *et al*, 2009; Wright *et al*, 2009). However, TLR2/4 deficient mice also lacking TLR9, which detects bacterial CpG-rich DNA, show reduced sensitivity to *Salmonella* and reduced intracellular bacterial replication. This is thought to be due to failures in endosome maturation that would normally trigger *Salmonella* to express SPI-2 (Talbot *et al*, 2009; Wright *et al*, 2009; Arpaia *et al*, 2011). In mouse infection models TLR4 activation has been shown to coordinate host survival and limit bacterial growth (Weiss *et al*, 2004).

The main PAMP recognised by TLR4 is LPS, which is found on the outer membrane of Gram-negative bacteria, such as *Salmonella*. LPS binds to TLR4 and its co-receptor MD2 (Park *et al*, 2009) with intracellular signalling orchestrated by homotypic interactions with the TLR4 Toll/interleukin-1 receptor (TIR)-domains and the TIR-containing adaptor proteins TIRAP and TRAM. These proteins form scaffolds for the binding of the TIR-containing MyD88 and TRIF respectively (Guven-Maiorov *et al*, 2015). TLR4 at the cell surface acts through the TIRAP-MyD88 pathway triggering many downstream signalling pathways including activating MAPK and NF-κB. The activation of MyD88-dependent signalling is shared with the other TLRs except for TLR3 (Gay *et al*, 2014; Kawasaki & Kawai, 2014; Guven-Maiorov *et al*, 2015). LPS stimulation also triggers TLR4 internalisation from the cell surface into endosomes (Tanimura *et al*, 2008; Zanoni *et al*, 2011; Ciesielska *et al*, 2020). Endosomal TLR4 signals through the TRAM/TRIF pathway leading to activation of alternative signalling pathways including IRF3 (Tanimura *et al*, 2008; Ciesielska *et al*, 2020). The TRIF-dependent signalling pathway is shared with TLR3 (Gay *et al*, 2014; Kawasaki & Kawai, 2014).

TLR4 stimulation leads to significant transcriptional and translational reprogramming as part of the inflammatory response (Kitamura *et al*, 2008; Schott *et al*, 2014). Activation of inflammatory transcription factors, including NF-κB and IRF3 lead to the transcription of immune response genes, including cytokines that signal to other immune cells. While the TLR4-dependent translational response is much less studied, through the use of purified LPS, it is known that translational upregulation of select mRNAs, including those encoding cytokines and negative feedback modulators of the inflammatory response, downstream of TLR4 activation is achieved by modulating the activity of RNA binding proteins that recognise cis-acting elements within these transcripts (Schott *et al*, 2014; Tiedje *et al*, 2012; Otsuka *et al*, 2019). In addition, changes in the translation of factors involved in mitochondrial oxidative phosphorylation have also been reported following TLR4 stimulation, leading to LPS-induced decreases in macrophage metabolism (Kitamura *et al*, 2008).

Rapid translational changes upon *Salmonella*-macrophage infection that are dependent on the SPI-1 injectisome have been previously identified (preprint: Wood *et al*, 2023). This includes SPI-1-dependent translational enhancement of genes whose expression is classically considered to be TLR4-dependent, such as pro-Il1β which showed clear SPI-1-dependent expression within 60 min (Ciesielska *et al*, 2020; Kelley *et al*, 2019; Bauernfeind *et al*, 2009; Mariathasan *et al*, 2004). *Salmonella* infection-dependent differences in translation could be detected as early as 5 min post-infection (preprint: Wood *et al*, 2023) and morphological changes in macrophages following TLR4 stimulation have also been observed in this timeframe (Kleveta *et al*, 2012). As such, the first hour of macrophage infection was investigated to identify TLR4-dependent changes in gene expression utilising knockouts of both host TLR4 and the bacterial SPI-1 machinery, demonstrating the ongoing host-pathogen battle whereby *Salmonella* supresses host protein production through transcriptional suppression, which is circumvented by rapid host translation response.

## Results

### TLR4 stimulation triggers rapid changes in macrophage morphology and cytokine production

*Salmonella* infection of macrophages is known to stimulate host actin polymerisation resulting in ruffling of the cell membrane (Jones *et al*, 1993; Perrett & Jepson, 2009). Both wildtype (WT) *Salmonella* and a strain unable to produce a functional SPI-1 injectisome, Δ*prgJ* (Man *et al*, 2014) - hereafter the injectisome mutant, induce ruffling in WT immortalised bone marrow derived macrophages (iBMDM) to similar levels within 15 minutes (Figure 1A, C). Macrophage lacking TLR4, on the other hand, exhibit ruffling when infected by WT *Salmonella* but not with *Salmonella* deficient in SPI-1 secretion (Δ*prgJ*). Stimulation with Pam3CSK4 (TLR2/1 agonist), flagellin (TLR5 ligand), or inert beads does not induce ruffling in either WT or TLR4-deficient macrophages, whereas LPS induces ruffling in a TLR4-dependent manner (Kleveta *et al*, 2012) (Figure 1). This suggests two independent mechanisms drive the rapid changes in macrophage morphology during *Salmonella* infection: an LPS-TLR4-dependent and a SPI-1 injectisome-dependent mechanism.

**Figure 1:**
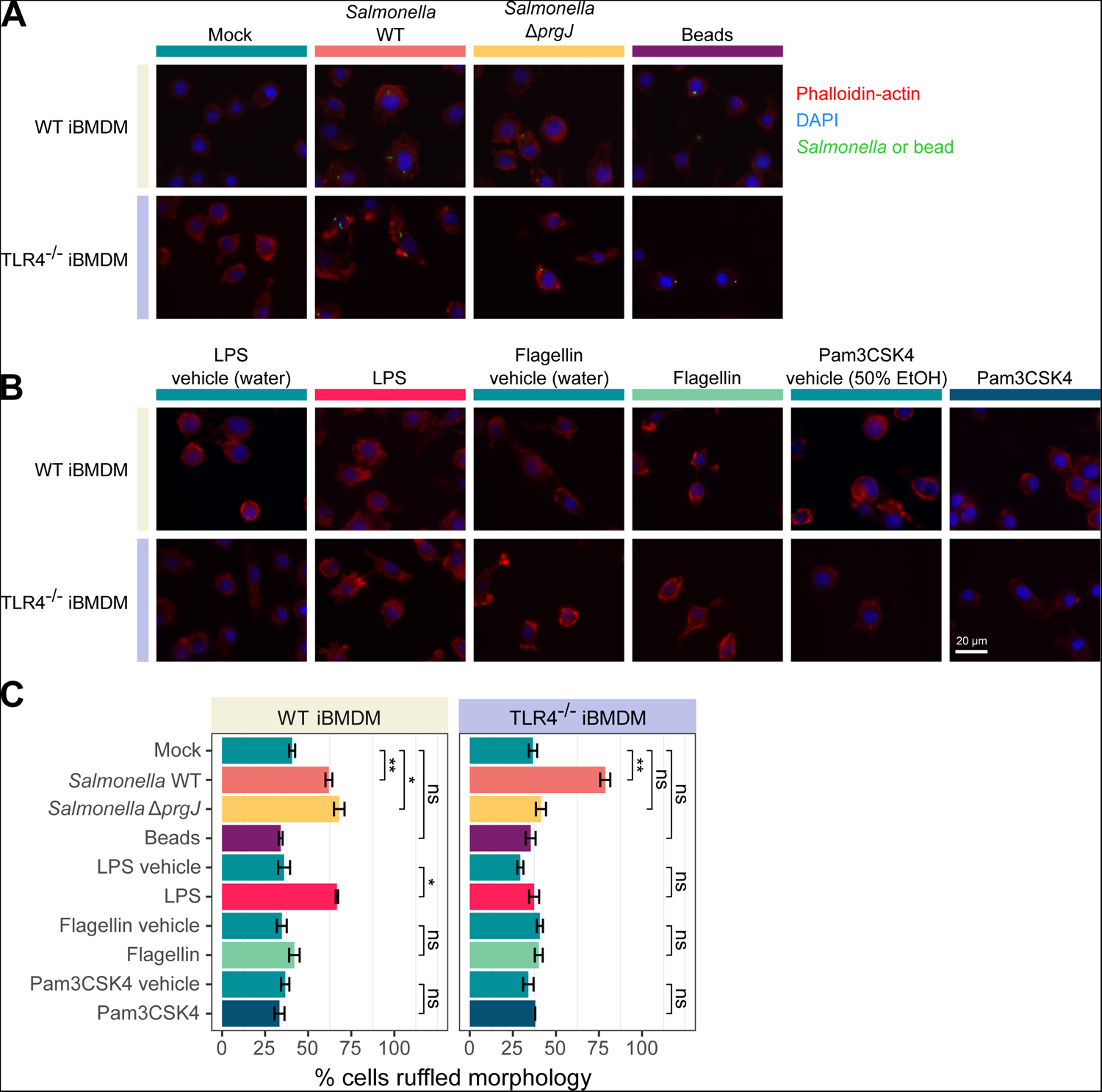
Infected macrophages undergo rapid TLR4 induced morphological changes during *Salmonella* infection. (**A** and **B**) Representative images of stimulated macrophages, 15 minutes post stimulation. Phalloidin-actin is shown in red; DAPI is shown in blue; and bacteria or beads are shown in green. (**A**) Cells stimulated with either WT or SPI-1 deficient, (Δ*prgJ*) *Salmonella* at MOI 10, or inert fluorescent beads at 100 beads per cell, or mock stimulated. (**B**) Cells stimulated with TLR agonists: 500 ng/ml LPS or 500 ng/ml Pam3CSK4 or 40 ng/ml flagellin, or the corresponding vehicle: water, 50% ethanol and water respectively. (**C**) Bar graph showing percentage of macrophages with ruffled morphology after 15 minutes of stimulation. P values determined by Holm adjusted one sided students t-test, ns indicates not significant, * indicates P value < 0.05 and ** indicates P value < 0.01. Error bars show standard error of the mean, n = 3.

Host cell death is characteristic of *Salmonella*-macrophage infection across mice and humans (Figure S1A) despite macrophages being a major host cell in systemic infection (Richter-Dahlfors *et al*, 1997; Monack *et al*, 1996; Hersh *et al*, 1999). Rapid macrophage death is attributed to the activation of the host inflammasome, triggering pyroptosis (Monack *et al*, 1996; Bryant, 2021; Hersh *et al*, 1999), and is independent of TLR4 activation (Figure S1). As most infected macrophages rapidly die, they will have a limited window in which they can produce proteins associated with innate immunity to coordinate the response to infection. Given this limited timeframe, we hypothesise targeted translational upregulation, which can be more rapid than transcriptional responses, plays an instrumental role in ensuring their robust production of proteins required for the innate immune response to *Salmonella* infection.

It is clear TLR4 stimulation rapidly alters the host cell (Figure 1) and drives much of the previously characterised innate response to Gram-negative bacteria, such as stimulating production of cytokines, including IL-1β (Figure S1B), as previously observed (Talbot *et al*, 2009). Further, TLR4 stimulation by LPS has been shown to increase the translational efficiency of transcripts encoding cytokines, such as *Tnf*, and other mRNAs containing AU-rich elements in their 3’ UTRs (Schott *et al*, 2014). However, the extent of rapid TLR4-dependent translational response, and whether this layer of regulation is actively perturbed by bacteria remains limited. As such, it is important to uncouple TLR4-dependent and independent gene expression responses within the context of *Salmonella* infection. In order to dissect the TLR4-dependent transcriptional response from the translational response, time-resolved parallel RNA-Seq and ribosome profiling (Ribo-Seq) was conducted on infected WT or TLR4^-/-^ primary BMDM, hereafter referred to as BMDM^WT^ and BMDM^TLR4^ respectively, with either WT or injectisome mutant *Salmonella* (Figure 2A).

**Figure 2:**
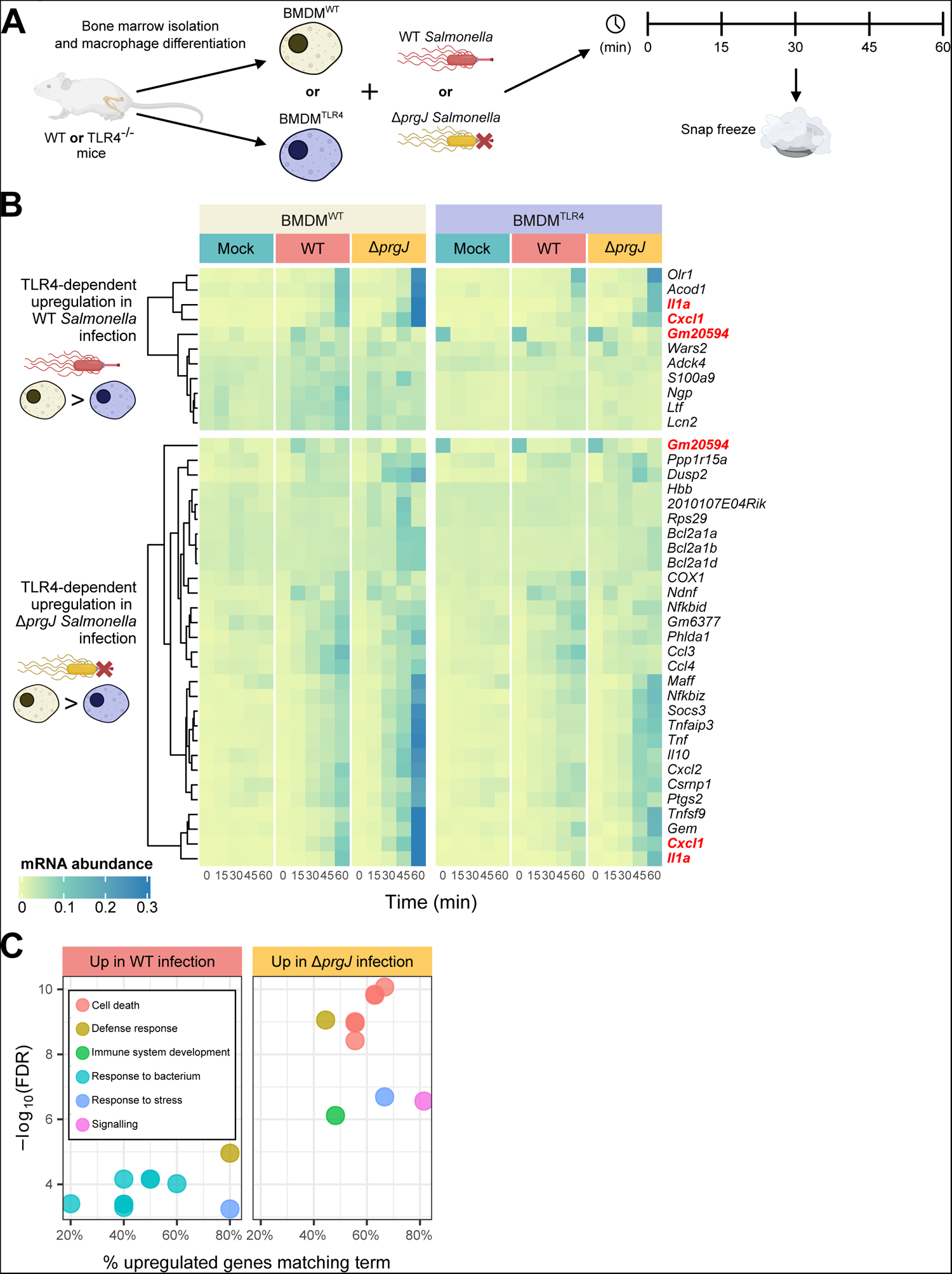
Transcripts upregulated on *Salmonella* infection of BMDM^WT^ over BMDM^TLR4^ are enriched for immune processes. Induction is greater when infected with *Salmonella* deficient in SPI-1 effector secretion. Data from two replicates. (**A**) *Salmonella* infected BMDM^WT^ or BMDM^TLR4^ were flash frozen every 15 min for the first hour of infection from which parallel RNA- and Ribo-Seq libraries were prepared. (**B**) Heatmaps showing relative RNA abundance of genes differentially expressed in (top) WT and (bottom) Δ*prgJ Salmonella* infection of BMDM^WT^ over BMDM^TLR4^ at two or more time points that are also differentially expressed in infection over mock infection of BMDM^WT^. Values are relative to total normalised RNA-Seq counts of that gene. Heatmaps use the same colour scale. Genes upregulated in both infections are highlighted in red. (**C**) Ten most significantly (smallest P value) enriched GO Biological Process terms of these differentially expressed genes, grouped by similarity. See Table S1 and S2 for the full list of significantly enriched GO terms. Figures produced with R and BioRender.

### Infected macrophages lacking TLR4 show reduced inflammatory transcriptional upregulation

Previous work has highlighted that the translational response precedes the transcriptional response in *Salmonella*-macrophage infection, with the translational response dominating over the first hour (preprint: Wood *et al*, 2023). Gene expression in the first hour of *Salmonella* infection was therefore examined to identify TLR4-dependent regulation (Figure 2A). While transcriptional responses are expected to be minimal within this timeframe, the few mRNAs whose abundance is rapidly upregulated were identified. Upregulation was defined as having log_2_ fold change (log_2_FC) greater than 1. The *Salmonella* injectisome and its effectors are known to influence the host transcriptional response (Galán, 2021; Sun *et al*, 2016; Jones *et al*, 2008; Pavlova *et al*, 2011; Jennings *et al*, 2017; Wang *et al*, 2020) and so, to identify the SPI-1-independent TLR4-dependent response, the infection was also conducted using the injectisome mutant *Salmonella*.

Initially, genes with upregulated transcript abundance in WT *Salmonella* infected BMDM^WT^ over BMDM^TLR4^ at any timepoint were selected, filtering out infection-independent differences by requiring them to also be upregulated at more than one timepoint in WT *Salmonella* infected over mock infected BMDM^WT^. This identified 11 genes with both TLR4- and *Salmonella* infection-dependent upregulation in the first hour of infection (Figure 2B, top panel). *Acod1*, *Olr1*, *Il1a* and *Cxcl1* showed a gradual increase in mRNA abundance over the first hour of both WT and injectisome mutant infection. In the absence of host TLR4, this increase is substantially reduced but some persists indicating the transcriptional upregulation is also driven by other pathways activated by *Salmonella*. The other genes whose transcript abundance was identified as TLR4-dependently upregulated by WT *Salmonella* infection show less change over the infection time course, indicating much of the difference in their abundance is not dependent on *Salmonella* infection.

*Acod1*, *Olr1*, *Il1a* and *Cxcl1* all showed greater increases when infected by the injectisome mutant *Salmonella*. As such, genes with TLR4-depenent upregulated transcript abundance in injectisome mutant *Salmonella* infection were also examined. Increases of a total of 29 genes were identified to be TLR4-dependent but injectisome independent across all time points. Of which, transcription of 13 genes (*Maff*, *Nfkbiz*, *Socs3*, *Tnfaip3*, *Tnf*, *Il10*, *Cxcl2*, *Csmp1*, *Ptgs2*, *Tnfsf9*, *Gem*, *Cxcl1* and *Il1a*) was greatest in injectisome mutant *Salmonella* infection (Figure 2B, bottom panel). While only 8% of identified genes were shared between WT and Δ*prgJ Salmonella* infection, the majority are upregulated in both but with a fold change less than 2 as used to identify them. Gene ontology (GO) term enrichment analysis of the genes showing TLR4-dependent upregulation in WT or injectisome mutant *Salmonella* infection reveals that, as expected, they have roles in immune system processes (Figure 2C; Table S1, S2). Enrichment analysis also highlighted the TLR4-dependent upregulation of genes involved in apoptosis in injectisome mutant *Salmonella* infection. This is driven by genes encoding BCL2A1 family proteins which are anti-apoptotic (Vogler, 2011). Interestingly, the upregulation of these genes was only seen in injectisome mutant *Salmonella* infection, where no induction of host cell death is observed in this timeframe (Figure S1).

### TLR4-induced changes in protein synthesis are not solely due to change in transcription

As described above, LPS stimulation has previously been shown through polysome profiling to alter the translation activity of macrophage-like cell lines such as J774A and RAW264.7 (Schott *et al*, 2014; Kitamura *et al*, 2008). This was also demonstrated in iBMDMs in this study (Figure S4A). Nevertheless, the stimulation of TLR4 during an infection cannot be fully recapitulated by purified LPS in solution alone. Additionally, detection of other PAMPs in infection may influence TLR4 signalling and bacterial effectors are known to modulate host immune signalling pathways (Galán, 2021; Sun *et al*, 2016; Jones *et al*, 2008; Pavlova *et al*, 2011; Jennings *et al*, 2017; Wang *et al*, 2020; Kawai & Akira, 2011; Schindler *et al*, 1990). As such, TLR4-dependent changes in protein synthesis during *Salmonella* infection were assessed through parallel ribosome profiling (Ribo-Seq) on the same samples as for RNA-Seq. Ribo-Seq is a highly sensitive method that reveals the global translatome at the time of harvest (Ingolia *et al*, 2009; preprint: Wood *et al*, 2023). The technique determines the position of ribosomes by exploiting the protection from nuclease digestion of a discrete fragment of mRNA (∼30 nucleotides) conferred by a translating ribosome. Deep sequencing of these ribosome-protected fragments (RPFs) generates a high-resolution view of the location and abundance of translating ribosomes on different mRNA species, reflecting the amount of synthesis of specific proteins. Due to the high-resolution data we obtained, we were able to accurately quantify protein synthesis by only considering in-frame reads that map to the coding sequence of transcripts, reflecting the movement ribosomes during the decoding process, prior downstream analysis. Genes with increased protein synthesis were similarly stratified as for RNA-Seq.

A total of 44 and 46 genes exhibited TLR4-dependent increases in protein synthesis in WT and injectisome mutant *Salmonella* infection respectively (Figure 3A-B). There was a greater overlap in genes showing TLR4-dependent increases in protein synthesis between infection with either WT or Δ*prgJ Salmonella* than in the RNA-Seq data, with 27% of genes identified as upregulated in both conditions. GO enrichment analysis of genes with increased protein synthesis identified similar roles to those with increased mRNA abundance: the response to bacteria, the defence response, with cell death specific to Δ*prgJ Salmonella* infection (Figure 3C). Significantly more genes were identified as upregulated in protein synthesis than RNA abundance, i.e. the increases in protein synthesis for upregulated genes cannot be explained by increases in RNA abundance alone (Figure 3D, S2, S3), highlighting their regulation at the level of translation.

**Figure 3:**
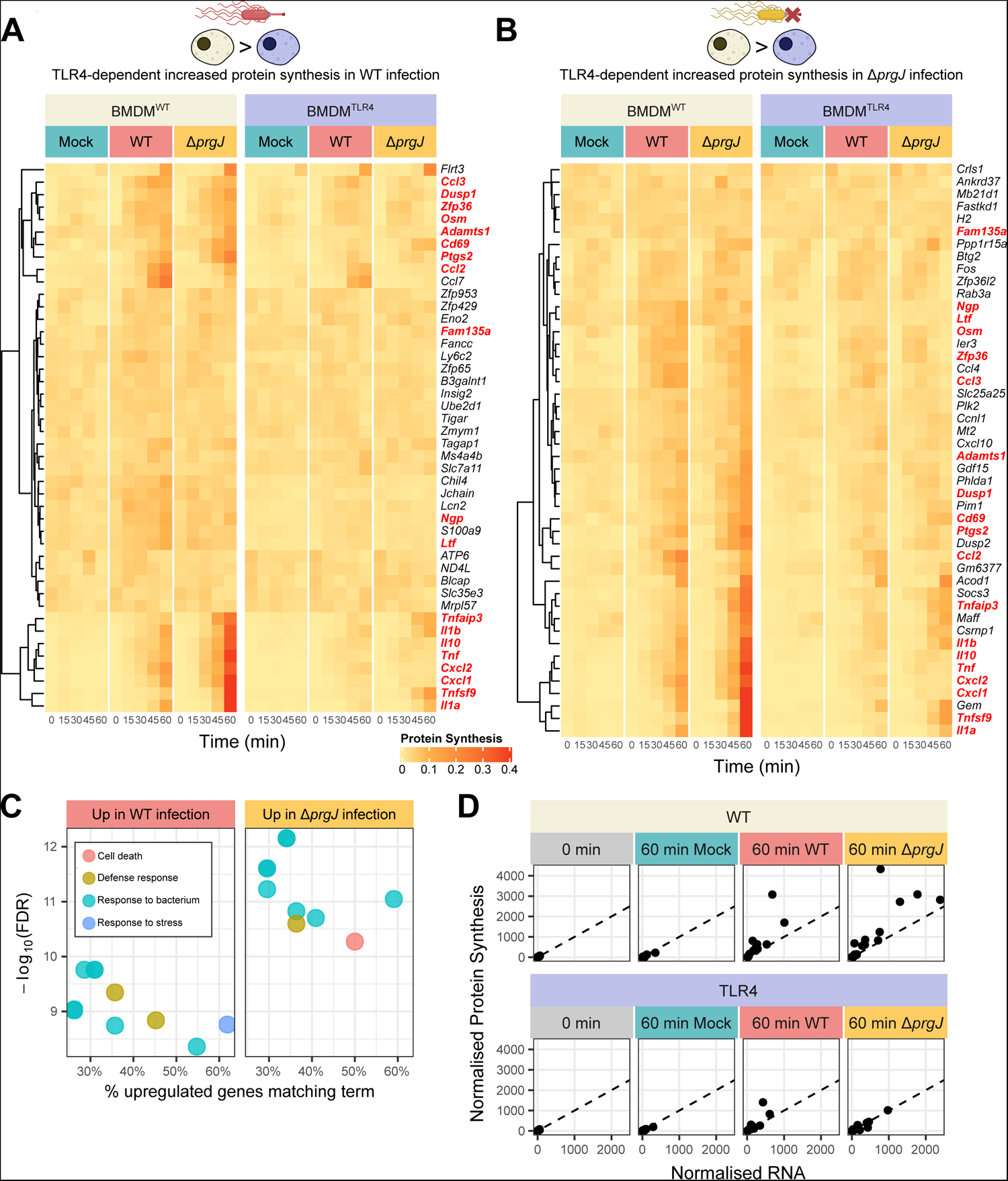
TLR4 dependent changes in protein synthesis are not always dictated by changes in mRNA abundance. Data from two replicates. (**A** and **B**) Heatmaps showing relative protein synthesis of genes differentially expressed in (**A**) WT and (**B**) Δ*prgJ Salmonella* infection of BMDM^WT^ over BMDM^TLR4^ at two or more time points that are also differentially expressed in infection over mock infection of BMDM^WT^. Values are relative to total normalised Ribo-Seq counts of that gene. Heatmaps use the same colour scale. Genes upregulated in both infections are highlighted in red. (**C**) Ten most significantly (smallest P value) enriched GO Biological Process terms of these differentially expressed genes, grouped by similarity. See Table S3 and S4 for the full list of significantly enriched GO terms. (**D**) Scatter plot comparing normalised RNA abundance and protein synthesis at 0 and 60 min post-infection for genes with TLR4-dependent upregulated protein synthesis in both WT and Δ*prgJ Salmonella* infection (red genes in **A** and **B**). Dashed line shows where RNA and protein synthesis are equal.

### TLR4-dependent translational upregulation of cell-signalling genes

The amount of protein produced will depend on both the abundance of its mRNA and the efficiency of their translation, which is independently regulated and occurs within the cytoplasm (Figure 4A). As such, translational efficiency (TE) was determined for all transcripts by comparison of the parallel RNA-Seq and Ribo-Seq data (Ingolia *et al*, 2009; preprint: Wood *et al*, 2023). Ribo-Seq captures the positions and abundance of translating ribosomes and so, in combination with RNA-Seq, can be used to determine the relative number of ribosomes per transcript, a measure of TE. Further, due to high proportion of highly phased reads in our data (Figure S4B), similarly to the quantification of protein synthesis, only in-frame reads are considered, i.e. reads that genuinely reflect ribosome occupation, therefore enable accurate quantification of TE for all downstream analysis.

**Figure 4:**
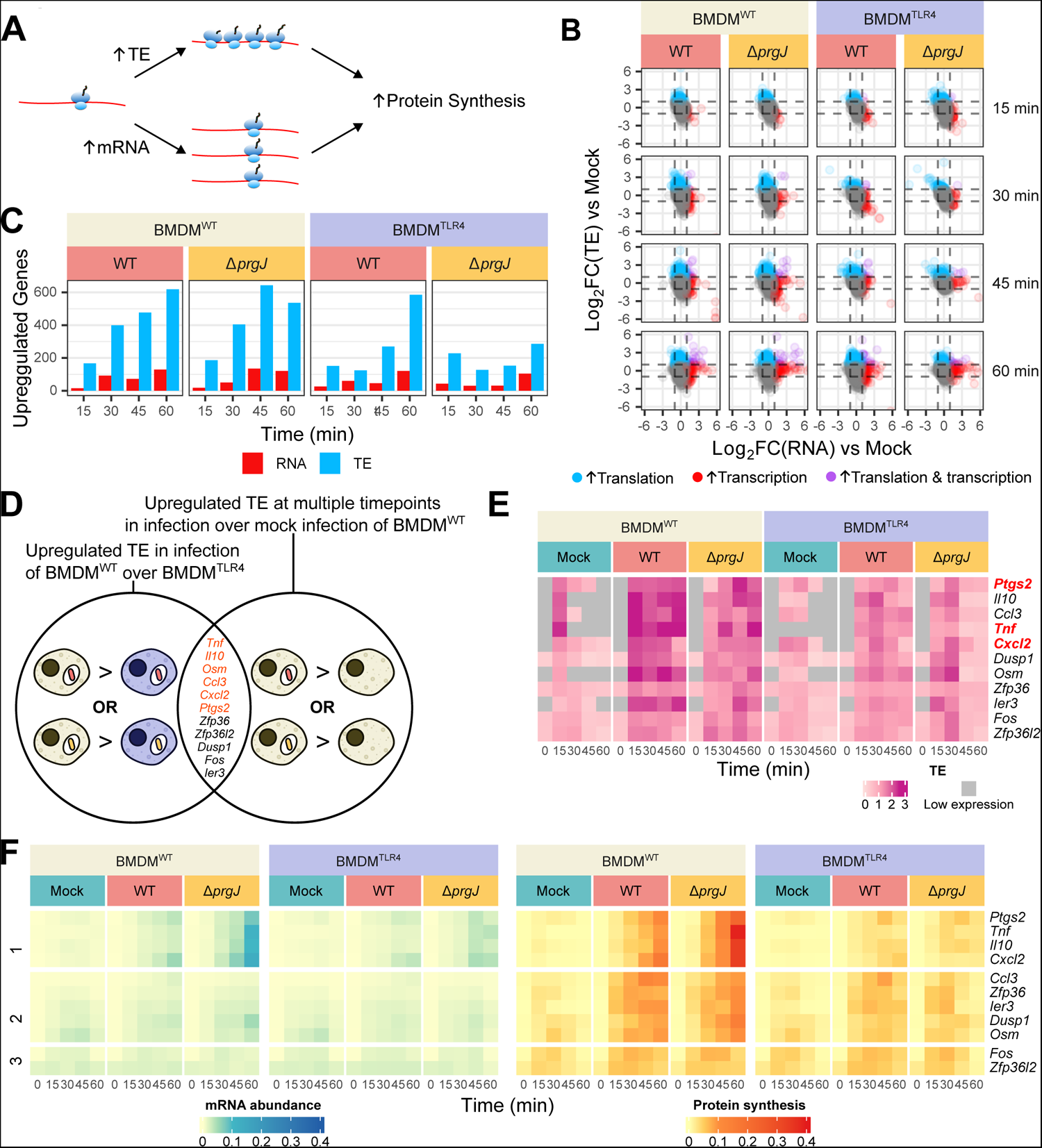
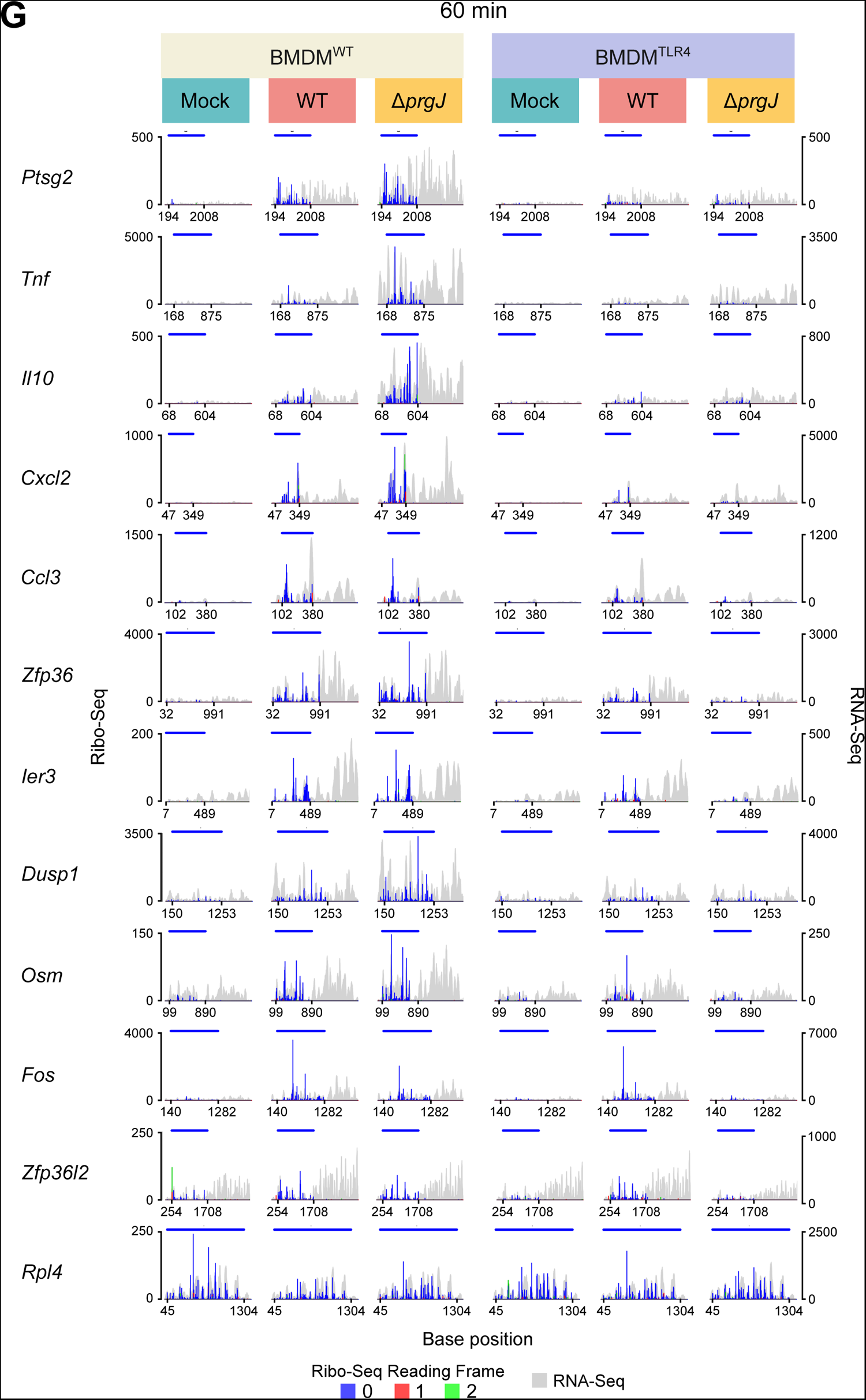
The TLR4-dependent, bacterial-effector-independent translational response. Data from two replicates. (**A**) Schematic showing two routes to increased synthesis of a particular protein: by increasing the number of its mRNAs or by increasing their translational efficiency. (**B**) Scatter plot comparing log_2_FC in RNA abundance and TE for the indicated condition vs mock infected host cells at the same timepoint. Dashed lines indicate log_2_FC vs mock of 1 and −1. Upregulated genes are coloured accordingly. (**C**) Bar plot showing number of genes upregulated transcriptionally or translationally. (**D**) Candidate selection criteria; genes were selected if they were upregulated at any timepoint between WT or Δ*prgJ Salmonella* infected BMDM^WT^ over BMDM^TLR4^ and were differentially expressed at 2 or more timepoints in infection of BMDM^WT^. Genes with direct roles in cell-cell signalling are coloured orange. (**E**) Heatmap showing TE of genes selected in C. An expression cut-off of 10 reads per samples was used for both RNA and protein synthesis, below which TE could not be calculated reliably. Genes in red were upregulated in WT *Salmonella* infection whereas all genes listed were upregulated in Δ*prgJ Salmonella* infection. (**F**) Normalised relative RNA- and Ribo-Seq counts, used to infer mRNA abundance and protein synthesis levels, respectively, for genes in **C**. Values are relative to total normalised RNA- and Ribo-Seq counts of that gene. K-means clustering (3 centres) of Ribo-Seq counts was used to group transcripts by translational behaviour throughout the infections. (**G**) Histogram showing inferred P site position of Ribo-Seq reads mapping to the indicated genes and coverage of RNA-Seq reads for samples at 60 min post infection. Ribo-Seq reads are coloured by their reading frame relative to the gene’s start codon.

Differential analysis of both TE and RNA abundance upon infection demonstrates variation in gene expression is predominantly driven by changes at the level of translation in the first 60 min of infection, with a greater number of genes upregulated translationally than transcriptionally in all conditions, aligning with previous reports (preprint: Wood *et al*, 2023) (Figure 4B, C). The combined absence of host TLR4 and bacterial SPI-1 T3SS substantially diminished the translational response.

Genes with significantly upregulated TE in BMDM^WT^ over BMDM^TLR4^ infected with either WT or injectisome mutant *Salmonella* were selected, filtered to select those also significantly upregulated in infection vs mock-infection of BMDM^WT^ at more than one timepoint (Figure 4D). Eleven genes with TLR4-dependent translational upregulation were identified, namely: *Ccl3*, *Cxcl2*, *Dusp1*, *Fos*, *Ier3*, *Il10*, *Osm*, *Ptgs2, Tnf*, *Zfp36* and *Zfp36l2* (Figure 4E, S3A). The fold-change between BMDM^WT^ and BMDM^TLR4^ was greatest in injectisome mutant infection (Figure S4C); however, WT *Salmonella* infection of WT cells gave rise to maximal TE (Figure 4E). This indicates that SPI-1 plays an important role in the translational upregulation of these genes, alongside TLR4.

Six of these genes showing TLR4-dependent translational upregulation are directly involved in cell-cell signalling, as cytokines and generators of signalling messengers, with the others also having key roles in modulating cell signalling pathways (Figure 4D). These genes were further divided into three clusters by their translational patterns throughout infection (Figure 4F): genes that exhibit rapid TLR4 and injectisome-dependent co-translational and transcriptional induction over time (Cluster 1); genes that exhibited rapid TLR4-dependent, injectisome independent translation induction with enhanced TE over time with little transcriptional response (Cluster 2); and genes that exhibit rapid TLR4 and injectisome-independent translational induction in response to infection with little transcriptional changes (Cluster 3). Due to the high resolution nature of the Ribo-Seq dataset, translation of these genes could be directly visualised at the sub-codon level (Figure 4G). Here, it is again clear that protein synthesis is not always linearly correlated with mRNA abundance and that while presence of *Salmonella* SPI-1 is able to limit TLR4-dependent transcriptional response, it does not repress the translational response, rather it likely acts synergistically to further increase TE of these genes.

## Discussion

Previous reports have demonstrated that both *Salmonella* (Jones *et al*, 1993; Perrett & Jepson, 2009) and purified LPS (Kleveta *et al*, 2012) can trigger actin ruffling in stimulated macrophages. Our results support these as two independent processes, i.e. the *Salmonella* trigger mechanism and LPS acting on TLR4, and demonstrates they both occur rapidly, within 15 min of exposure. Activation of other cell surface TLRs did not result in an increase in ruffling, consistent with previous reports that LPS-induced macrophage actin ruffling is a MyD88-independent process (Kong & Ge, 2008; Wall *et al*, 2017).

The transcriptional response within the 15-60 min of *Salmonella* infection is, as expected, limited. The few genes identified as differentially transcriptionally regulated are associated with immune processes, in alignment with previous reports of the transcriptional response to LPS stimulation of TLR4 (Ciesielska *et al*, 2020; Kelley *et al*, 2019). In contrast, a greater translational response was observed in both WT and SPI-1-deficient *Salmonella* infection than transcriptional response, due to translational modulation of pre-existing mRNAs in the cytoplasm and aligns with our previous study on *Salmonella* induced translational regulation (preprint: Wood *et al*, 2023). Surprisingly, much of this rapid translational response was lost in the absence of both bacterial SPI-1 injectisome and host TLR4, suggesting their predominant role in driving translational responses within the first hour of *Salmonella* infection of macrophages.

More genes were identified as TLR4-dependently transcriptionally upregulated in infection with SPI-1-deficient *Salmonella* than WT. This was also true for genes with enhanced protein synthesis. Many of these upregulated genes still showed upregulation in WT *Salmonella* infection, though to a lesser degree than in injectisome mutant *Salmonella* infection. Changes in transcription and changes in protein synthesis were found to be non-linearly related, highlighting the role of altered TE in regulating expression from a given transcript. Genes identified with significant TLR4-dependent upregulation of TE were mostly involved in cell-to-cell signalling. Interestingly, the TE of these genes in WT *Salmonella* infection exceeded that in injectisome mutant *Salmonella* infection, despite lower transcript abundance. It is therefore clear that the *Salmonella* SPI-1 injectisome plays an important role in attenuating the TLR4-dependent transcriptional response, likely due to SPI-1 effector-dependent modulation of signalling downstream of TLR4 that has previously been described (Galán, 2021; Sun *et al*, 2016; Jones *et al*, 2008; Pavlova *et al*, 2011; Jennings *et al*, 2017; Wang *et al*, 2020), but the translational response is not inhibited in this way. Rather, the translational response is maximal with intact SPI-1 injectisome, suggesting that it in some way augments the TLR4-dependent increases in translation of these transcripts. This may be due to the action of effectors secreted into the host cell through the SPI-1 injectisome or as a response to injectisome mediated host membrane injury as has previously been demonstrated for other translational responses (preprint: Wood *et al*, 2023).

The swift synthesis of cytokines is of particular importance in *Salmonella* infection given the rapid induction of SPI-1-dependent host cell death, leaving only a short window for most infected macrophages to mount an immune response. Modulation of the translational efficiency of cytokine encoding mRNAs in parallel with increasing their abundance allows for a more robust response. This is particularly important where the transcriptional arm of the host response is antagonised by bacterial effectors.

## Acknowledgements

G.W. was supported by the Department of Pathology PhD studentship and F.L. was supported by a BBSRC DTP studentship. B.Y.W.C., R.J., G.W. F.L. and M.B. supported by a Medical Research Council Fellowship and BBSRC project grants to B.Y.W.C. [MR/R021821/1, BB/X001261/1, BB/V017780/1 and BB/V006096/1]

## Author contributions

B.Y.W.C., and C.B. conceived the research. B.Y.W.C., C.B., G.W., and R.J., designed experiments. R.J. and G.W. performed molecular and cell biology experiments, R.J., M.B., G.W., and F.L. performed bioinformatics, P.T. maintained mice colonies as well as mice processing. B.Y.W.C., and G.W. wrote the manuscript.

## Methods

### Salmonella infection

Primary BMDMs were isolated and differentiated as previously described (Man *et al*, 2014) from WT C57BL/6 mice or TLR4^-/-^ C57BL/6 mice. iBMDM were generated by viral transformation of BMDMs from these mice as previously described (De Nardo *et al*, 2018). All cells were routinely grown in DMEM supplemented with 10% foetal bovine serum at 37°C, 5% CO_2_. Media for primary BMDM was supplemented with 20 ng/ml M-CSF (Peprotech). Cells were infected at a multiplicity of infection of 10 with WT or Δ*prgJ S*. Typhimurium SL1344 as previously described (preprint: Wood *et al*, 2023). Briefly, bacteria were grown to late exponential phase, washed in cell culture media, and added to the cultured cells. At 15 min post infection, gentamicin (50 µg/ml final concentration for primary BMDM, 100 µg/ml final concentration for all others) was added to ensure synchronous infection.

### Cell stimulation and immunofluorescence microscopy

Cells were plated onto glass coverslips and grown until 50% confluency before being infected or stimulated as indicated. 1 µm Fluoresbrite Yellow Green Microspheres (Polysciences) were washed in cell culture media and used as a control for phagocytosis at 100 beads per cell. TLR4 was stimulated with *Escherichia coli* O127:B8 LPS (Sigma-Aldrich) at 500 ng/ml. TLR5 was stimulated with recombinant *S*. Typhimurium flagellin (Abcam) at 40 ng/ml. TLR1/2 was stimulated with Pam3CSK4 (Tocris). Mock stimulations were conducted with vehicle only: water for LPS and flagellin; 50% ethanol for PAM3CSK4; media only for beads and infections.

Cells were fixed with 4% paraformaldehyde and stained with Phalloidin-CF594 conjugate (Biotium), DAPI (Invitrogen), and goat anti-CSA (Insight Biotechnology) followed by donkey anti-goat IgG conjugated to Alexa-488 (Abcam). Cells were mounted onto slides and imaged at 100x magnification. 20 random fields were captured per replicate and cells were classified as ruffled or not as described previously (Kleveta *et al*, 2012). 50 cells were chosen at random to represent each replicate. One-sided student’s t-test with Holm correction was used to identify significant increases in ruffling compared to the corresponding mock or vehicle stimulation.

### Ribosome profiling and RNA sequencing

Combined ribosome profiling and RNA sequencing of infected macrophages was conducted as previously described (preprint: Wood *et al*, 2023). Briefly, culture supernatant was removed from cells, and they were flash frozen. Cells were lysed in buffer containing cycloheximide and chloramphenicol, and lysates were split for parallel RNA-Seq and Ribo-Seq. For Ribo-Seq, lysates were treated with RNase I and fragments protected from digestion by the ribosome were purified. For RNA-Seq, total cellular RNA was fragmented by alkaline hydrolysis. This was followed by library generation as previously described (Irigoyen *et al*, 2016; Chung *et al*, 2015, 2017). Sequencing was performed using NextSeq-500 (Illumina).

### Bioinformatics

Reads from two replicate infections were combined and aligned sequentially to mouse rRNA and mouse mRNA. Mouse reference sequences were based on release mm10. Counting and differential expression in both RNA abundance and translational efficiency was performed using riboSeqR (Chung *et al*, 2015) and Xtail (Xiao *et al*, 2016). GO term enrichment was performed using g:Profiler (Peterson *et al*, 2020). Data analysis was performed and figures were generated using R. Diagrams were created with BioRender.com.

### Cytotoxicity assays

Cells were plated in 24-well plates and grown until 70-80% confluency before being infected as indicated. Cytotoxicity was measured by lactate dehydrogenase (LDH) release using CytoTox96 Non-radioactive Cytotoxicity Assay kit (Promega) according to the manufacturer’s instructions. Values are relative to total host lysis in 0.9% Triton X-100.

### Enzyme-linked immunosorbent assay (ELISA)

Culture supernatants were removed from infected cells at the indicated timepoint. ELISAs were performed to quantify IL-1β in these culture supernatants using the Mouse IL-1 beta/IL-1F2 DuoSet ELISA kit (R&D Systems) per the manufacturer’s instructions.

### Polysome profiling

Polysome profiling has been described previously (Chassé *et al*, 2017). Briefly, cell lysates were prepared as for ribosome profiling and then centrifuged over a linear 10-45% sucrose gradient prepared using a Gradient Master (BioComp). RNA concentration, measured by absorbance at 254 nm, was then traced across the gradient using a Density Gradient Fractionation System (Brandel).

