## Supplemental figures for "Activity of *Salmonella* SPI-1 inhibits the TLR4-dependent transcriptional but not translational response during macrophage infection"

### Figure S1

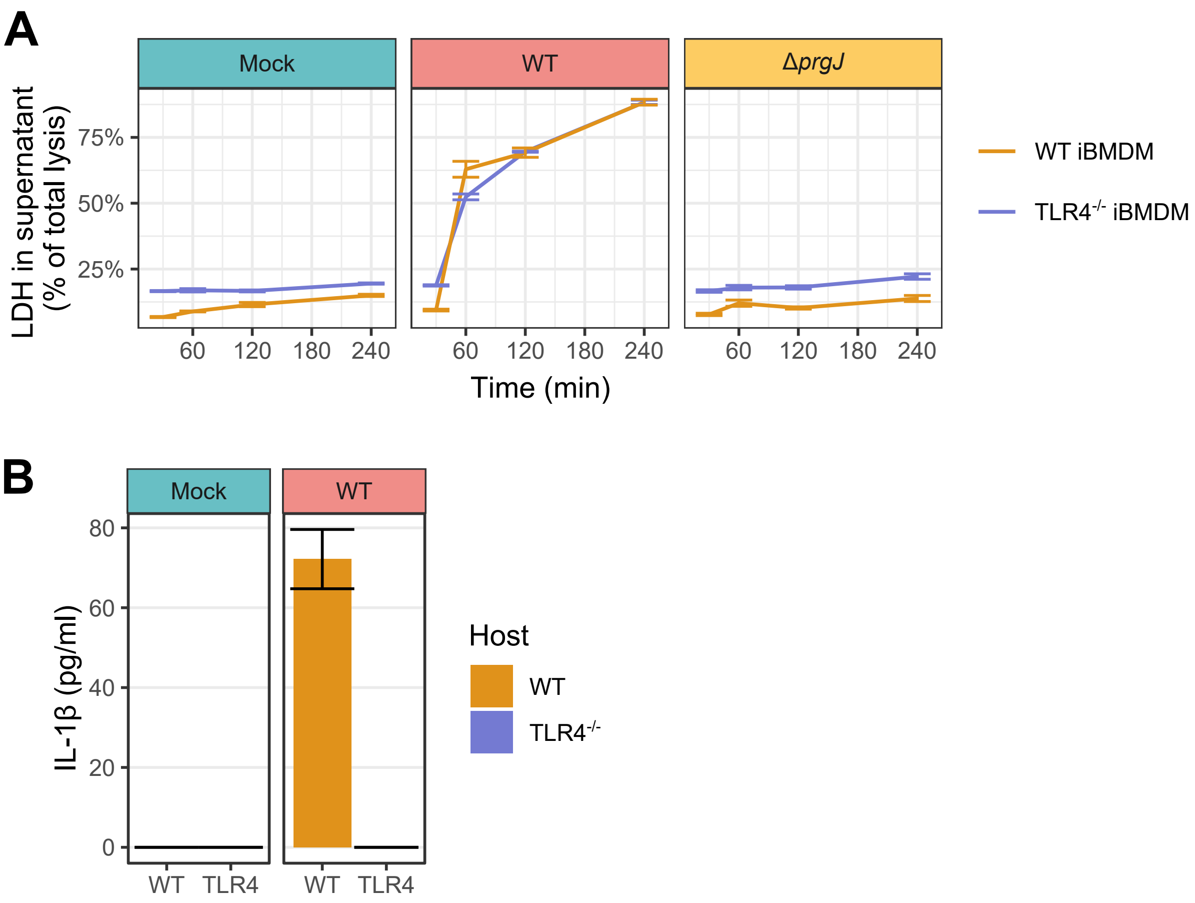


***Figure S1:*** (**A**) Cytotoxicity of WT or Δ*prgJ Salmonella* infection in WT and TLR4$^{-/-}$ iBMDMs as measured by LDH release into culture supernatant. Values are relative to those following total cell lysis. N=2, data representative of 2 independent experiments. (***B***) IL-1β in culture supernatant 4 hours after WT *Salmonella* infection of WT and TLR4$^{-/-}$ iBMDMs as determined by ELISA. N=3.

### Figure S2

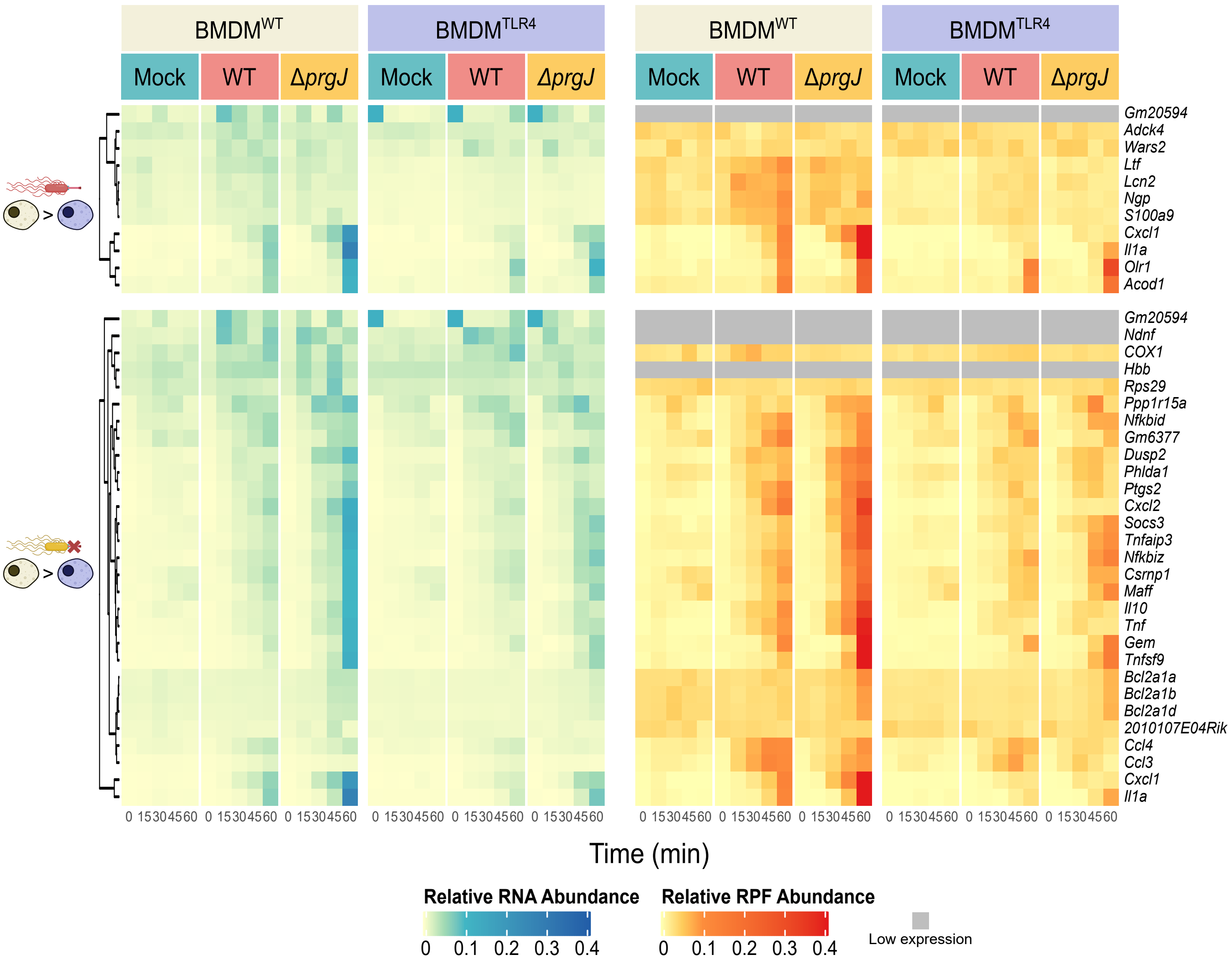


**Figure S2**: Expression of genes transcriptionally upregulated in a TLR4-dependent manner over *Salmonella* infection (WT: top, Δ*prgJ*: bottom) at the levels of both transcript abundance and protein synthesis. Genes are clustered by RNA abundance. Values are relative to total normalised RNA- and Ribo-Seq counts of that gene.

### Figure S3

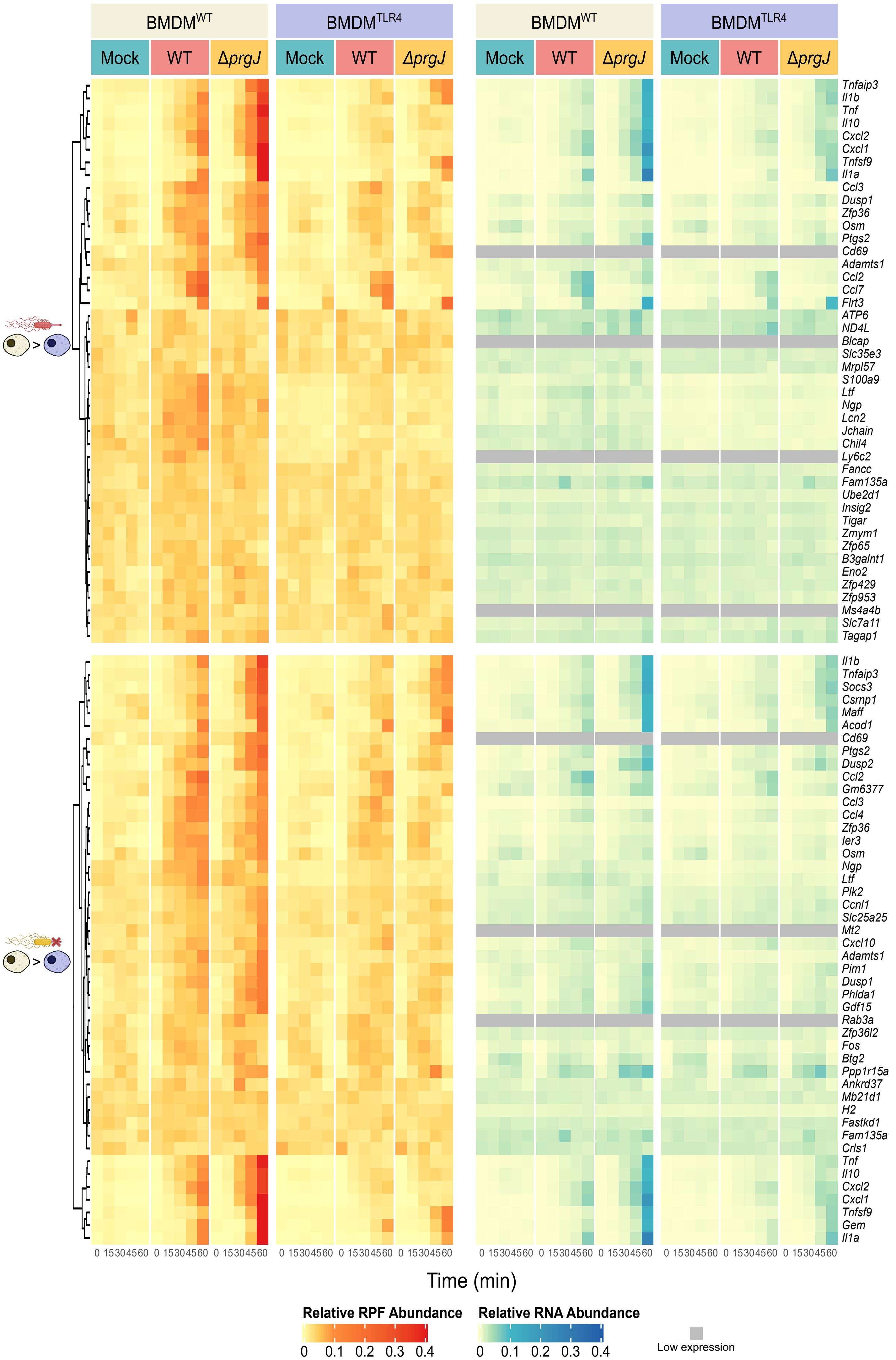


**Figure S3**: Expression of genes with upregulated protein synthesis in a TLR4-dependent manner over *Salmonella* infection (WT: top, Δ*prgJ*: bottom) at the levels of both transcript abundance and protein synthesis. Genes are clustered by amount of protein synthesis. Values are relative to total normalised RNA- and Ribo-Seq counts of that gene.

### Figure S4

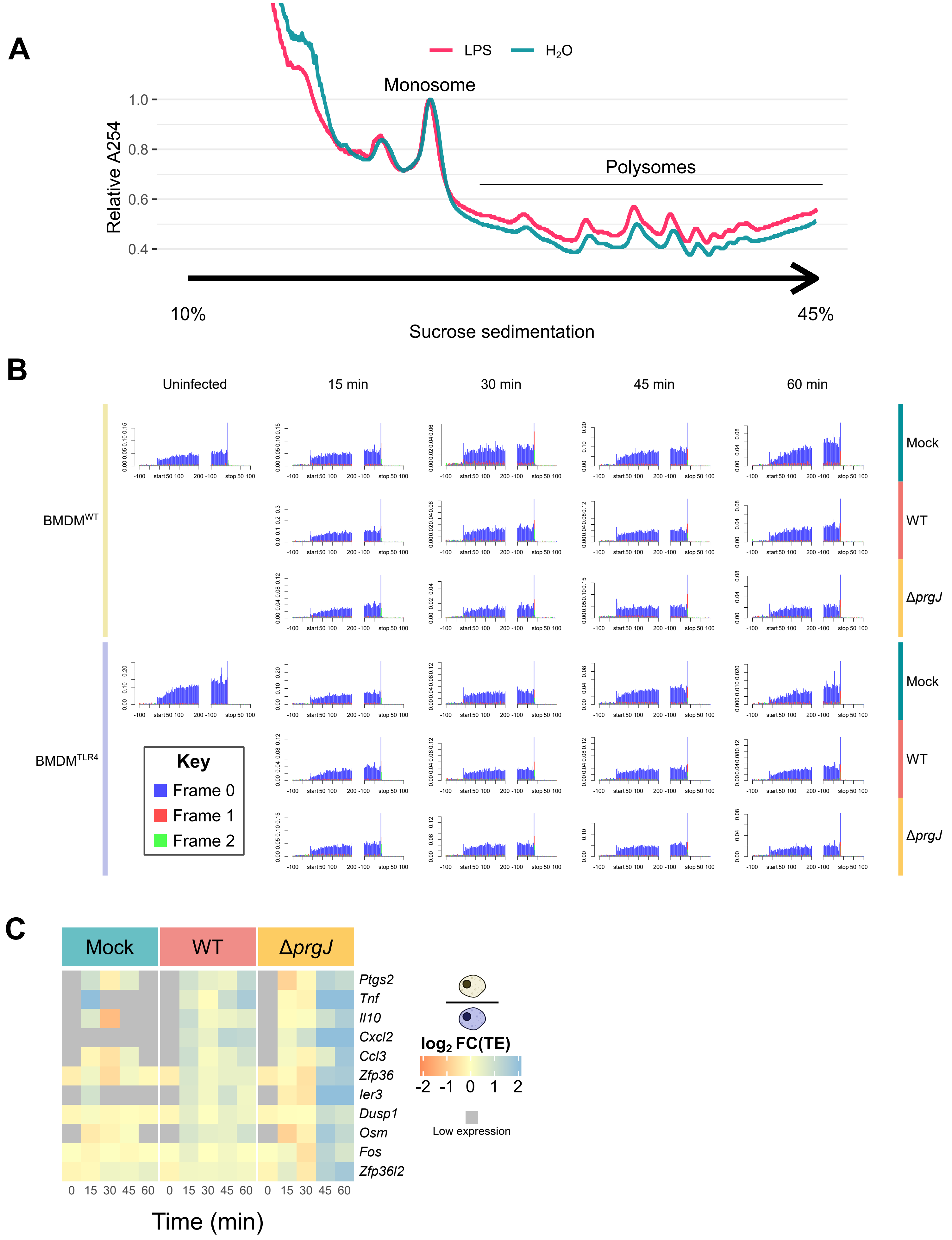


**Figure S4**: (**A**) Polysome profile trace of WT iBMDMs stimulated with 500 ng/ml LPS or water vehicle for 60 min. Traces are scaled to equalise their monosome peak. Representative of two independent experiments. (***B***) Metagene translatome from ribosome profiling of Salmonella infected BMDM$^{WT}$ and BMDM$^{TLR4}$ over the first 60 min of infection. Histograms show ribosome protected fragment 5′ ends relative to start and stop codons coloured by their reading frame relative to the first codon of the coding sequence. (***C***) Heatmap showing the $\log_2$FC of translation efficiency (TE) in infected BMDM$^{WT}$ over BMDM$^{TLR4}$ for genes with significantly upregulated TE. An expression cut off of 10 reads was used for both RNA and protein synthesis, below which TE could not be calculated reliably.
